# Reliable Molecular Retrieval from Mass Spectra using Conformal Prediction

**DOI:** 10.64898/2026.03.12.711424

**Authors:** Morteza Rakhshaninejad, Gaetan De Waele, Mira Jürgens, Willem Waegeman

## Abstract

A key task in the computational analysis of liquid chromatography–tandem mass spectrometry (LC– MS/MS) data is identifying the molecular structure underlying a measured spectrum. A common approach ranks candidate molecules retrieved from chemical databases using predicted fingerprint similarities, yet standard metrics such as top-*k* accuracy summarize performance only at the dataset level and provide no spectrum-specific reliability statement. In this work, we apply conformal prediction to candidate-based molecular retrieval to construct spectrum-specific prediction sets that contain the true molecule with a user-specified probability. We evaluate marginal and conditional conformal prediction across three experimental scenarios representing in-distribution, partially shifted, and fully out-of-distribution settings on the MassSpecGym benchmark. When calibration and test data are aligned, conformal prediction attains the target coverage with small candidate sets for most spectra. Under distribution shift, prediction sets become larger as rankings grow more ambiguous, although candidates can still be reduced when calibration remains representative. Conditional conformal prediction improves subgroup reliability across spectra of different difficulty, with the best gains obtained using confidence-based grouping. Overall, conformal prediction turns candidate rankings into reliable, spectrum-specific candidate sets with an explicit reliability–efficiency trade-off.

## 1 Introduction

Liquid chromatography–tandem mass spectrometry (LC–MS/MS) is a widely used technique for detecting and identifying small molecules in metabolomics.^1–3^ Recent advances in machine learning, including deep learning models trained on MS/MS data, have substantially improved spectrum representations and downstream identification performance.^4,5^ At the same time, mapping an MS/MS spectrum to a molecular structure remains challenging.^6^ Broadly speaking, three categories of methods exist:^7^ (1) spectral library search, which restricts annotation to molecules for which experimentally derived spectra are available; (2) molecular retrieval, which expands the search space to molecules included in chemical databases; and (3) *de novo* molecule generation, which can in principle predict novel structures. Since *de novo* methods remain largely ineffective for chemically novel compounds under realistic conditions of structural diversity,^1,8^ molecular retrieval offers a practical balance between chemical coverage and accuracy. The paradigmatic approach to molecular retrieval proceeds by predicting a fingerprint or embedding that can be matched against large chemical databases to retrieve and rank candidate molecular structures based on similarity scores.^9,10^ In practice, the search space is enormous: chemical databases can contain tens to hundreds of millions of molecules, making exhaustive ranking computationally infeasible. To address this, candidate-based identification pipelines restrict the search to a manageable subset, most commonly using precursor mass or molecular formula constraints.^11^

Despite candidate filtering, reliable identification from LC–MS/MS remains challenging. Even after restricting candidates by mass or formula, candidate sets can be large and vary widely in both size and how easily candidates can be distinguished from one another. This variation arises because distinct molecules can generate similar fragmentation patterns. As a result, some spectra yield confident rankings over only a few plausible candidates, while others yield highly ambiguous rankings across much larger candidate spaces.^6^

Standard evaluation metrics such as top-*k* accuracy quantify retrieval performance at the dataset level by checking whether the correct molecular structure appears among the top-*k* ranked candidates.^1^ For example, top-1 accuracy is the proportion of spectra for which the correct molecule is ranked first. While useful for benchmarking, top-*k* accuracy does not quantify reliability at the level of an individual spectrum. It does not tell us how many candidates should be retained for a given spectrum so that the true molecule is included with high probability, nor does it reflect how confidence varies across spectra with different candidate-set sizes or score separations. In many cases, the model assigns a clearly dominant score to the top candidate and only a small subset would be sufficient, whereas in more ambiguous cases several candidates can receive similar scores and a larger subset is needed. This motivates set-valued outputs that return, for each spectrum, a spectrum-specific subset of candidates that can be small in easy cases but expands when the spectrum is intrinsically ambiguous, together with an explicit reliability statement.

Conformal prediction (CP) provides a principled approach for constructing such set-valued outputs with finite-sample coverage guarantees.^12^ Given a user-specified confidence level 1 − *α* (e.g., 90%), CP constructs, for each spectrum *x*, a prediction set 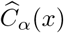 such that, under exchangeability, the true label is contained with probability at least 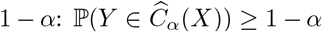.^13^ CP has recently found applications in cheminformatics ranging from ADMET property prediction,^14^ toxicity modeling,^15,16^ and efficient compound screening^17^ to modeling cancer cell growth inhibition under strong distribution shifts^18^ and antimicrobial resistance detection from mass spectra,^19^ underscoring its growing relevance to analytical chemistry workflows. In the context of LC–MS/MS candidate-based retrieval, CP can be used to output a prediction set of candidate molecules that is typically a subset of the full candidate set and adapts in size to the difficulty of each spectrum. However, the guarantees provided by standard conformal prediction are marginal and therefore hold only on average over the test distribution.^20^ Achieving marginal coverage does not imply that coverage is stable across meaningful subsets of spectra. LC–MS/MS retrieval is inherently heterogeneous, and several characteristics vary strongly from one spectrum to another, including the size of the candidate set, how concentrated the model scores are, the score gap between the top-ranked candidates, the precursor mass, and other spectrum or molecular characteristics. These factors are directly related to retrieval difficulty. As a result, a method can meet the target marginal coverage while still producing systematic coverage disparities across sub-populations, for instance by under-covering spectra with large candidate sets or highly ambiguous rankings and over-covering spectra with small candidate sets and clear score separation. This behavior is problematic in metabolomics applications, where practitioners need uncertainty estimates that remain reliable for the specific types of spectra they encounter, not only on average across an entire benchmark.

These considerations motivate conditional CP for candidate-based molecular retrieval.^21^ Instead of controlling coverage only on average, the objective is to obtain coverage that remains stable across meaningful sub-populations defined by spectrum characteristics, candidate-set properties, and model-derived uncertainty signals. This is particularly relevant in LC–MS/MS retrieval because the difficulty can vary drastically between spectra, and users often face the hardest cases in practice, such as large candidate spaces with weak score separation. Achieving conditional coverage in stronger distribution shift retrieval settings is challenging for several reasons. Relevant conditioning variables are often continuous and can be strongly correlated, and several reasonable choices may capture overlapping aspects of retrieval uncertainty. As a result, it is not always clear which variables define distinct subgroups of spectra. In addition, the relationship between these variables and retrieval difficulty can change across experimental scenarios and data splits, especially under distribution shift when candidate generation and spectrum distributions differ between calibration and test data.^22,23^ Finally, enforcing conditional guarantees can increase prediction set sizes, which exposes an inherent trade-off between reliability and efficiency in downstream analysis. Understanding which conditioning strategies are meaningful, how conditioning variables interact, and how conditional CP methods behave across stronger distribution shift scenarios is therefore essential for building uncertainty-aware molecular identification systems that are both reliable and practical.

In this work, we study both marginal and conditional CP for LC–MS/MS–based candidate retrieval. We focus on candidate-based identification settings in which a spectrum is associated with a predefined candidate set, and the goal is to output a smaller prediction set of candidates with a user-specified coverage target. To address heterogeneity across spectra, we investigate conditioning strategies that form sub-populations using clustering^24^ and nearest-neighbor groupings^25,26^ based on spectrum, candidate-set, and model-derived characteristics. We evaluate these approaches across multiple experimental scenarios that represent increasingly challenging settings and include distribution shift between calibration and test data. Our main contributions are summarized as follows.

- Multiple non-conformity scores, including adaptive and regularized variants, are compared by coverage and set size.
- Three scenarios of increasing difficulty range from independent and identically distributed (IID) to out-of-distribution (OOD), evaluating conformal prediction outputs.
- Correlation analysis of candidate-level and model-derived variables is used to select informative conditioning variables.
- Clustering-based and nearest-neighbor conditional CP methods are evaluated, quantifying the reliability– efficiency trade-off under shift.

The remainder of the paper is organized as follows. Section 2 presents the methodological framework. We first describe the candidate-based retrieval setting, the construction of prediction sets, conformal prediction, and the evaluation metrics. We then introduce the non-conformity scores and the conditional conformal approaches considered, including clustering-based and nearest-neighbor constructions and the conditioning variables used to define sub-populations. Finally, we summarize the experimental setup, including the dataset, experimental scenarios, retrieval model, and conformal prediction configuration. Section 3 reports empirical results, first characterizing retrieval difficulty across scenarios and then analyzing marginal and conditional conformal performance, including subgroup behavior and the reliability–efficiency trade-off. Finally, Section 4 summarizes the main findings, discusses limitations, and outlines directions for future work.

## 2 Methods

### 2.1 Candidate-based molecular retrieval

In candidate-based LC–MS/MS retrieval, each spectrum is represented by a feature vector 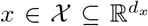. Let 𝒴 ⊆ {0, 1}^*D*^ denote the space of molecular fingerprints, where *D* ∈ ℕ is the fingerprint dimensionality. For each spectrum *x*, the goal is to identify the corresponding fingerprint *y* ∈ 𝒴. Candidate-based pipelines associate each spectrum *x* with a finite candidate set *𝒜* (*x*) ⊆ 𝒴 obtained through external constraints, most commonly precursor mass and optionally molecular formula. ^**27**^ The evaluation setting assumes that the true fingerprint belongs to the candidate set, i.e., *y* ∈ 𝒜 (*x*). In fingerprint-based retrieval pipelines, the spectrum is mapped to a predicted fingerprint (or embedding) 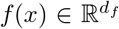, where *d*_*f*_ = *D* in the experiments. Each candidate *c* ∈𝒜 (*x*) is represented by a fingerprint vector *ϕ*(*c*) ∈ {0, 1}^*D*^. A retrieval model assigns each candidate *c* ∈ 𝒜 (*x*) a score *s*(*x, c*) ∈ ℝ, where larger scores indicate higher plausibility. In the fingerprint-based setup, the score is computed by comparing the predicted fingerprint *f* (*x*) to the candidate fingerprint *ϕ*(*c*) via a similarity function,

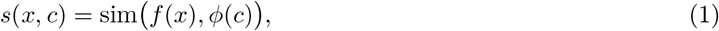

where sim(·, ·) denotes the similarity used for candidate scoring in the retrieval pipeline (cosine similarity in our experiments). Candidates are ranked by decreasing score, yielding an ordered list (*c*_(1)_, *c*_(2)_, …, *c*_(|𝒜 (*x*)|)_) such that

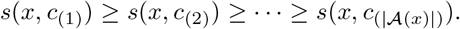

The ranked list induced by *s*(*x, c*) serves as the basis for constructing set-valued outputs with reliability guarantees.^9^ In the experiments, *f* (·) corresponds to a multi-layer perceptron (MLP) that is trained to optimize candidate ranking over 𝒜 (*x*) using a listwise rank-based cross-entropy objective with temperature scaling and dropout.^28^ At inference time, candidates are scored using Equation (1). These scores are then converted into probabilities *π*(*x, c*) via a softmax operation over 𝒜 (*x*) and are mapped to non-conformity scores *r*(*x, c*) as described in Section 2.3. For each spectrum *x*, the objective is to output a prediction set 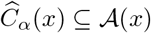 that contains the true fingerprint with high probability while remaining as small as possible. Let (*X, Y*) denote a random test pair drawn from the evaluation distribution. This objective can be written as minimizing the expected set size subject to a coverage constraint at error level *α* ∈ (0, 1):

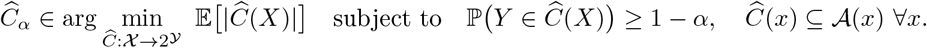

The size of 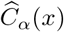 is allowed to vary across spectra, so that spectra with clear score separation lead to small prediction sets while ambiguous spectra can produce larger ones. Prediction sets are evaluated through coverage, defined as the frequency with which the true fingerprint *y* is included in 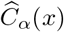, and efficiency, quantified by the prediction set size 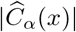 or, since candidate set sizes |𝒜 (*x*)| can vary substantially across spectra, by the relative size 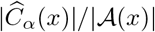.

### 2.2 Conformal Prediction

Conformal prediction (CP) provides a general framework for constructing prediction sets with finite-sample coverage guarantees under an exchangeability assumption.^12,13^ A calibration set 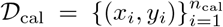 is assumed to be available, and the calibration pairs are assumed to be exchangeable with the test samples 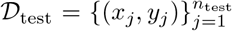. Under this assumption, CP uses the calibration data to translate model scores into prediction sets 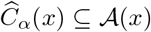 whose inclusion of the true molecule can be guaranteed at a user-specified error level *α* ∈ (0, 1). Specifically, for a random test pair (*X, Y*), conformal prediction ensures the finite-sample guarantee 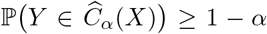. Since LC–MS/MS benchmarks can involve distribution shifts between calibration and test data, robustness of conformal procedures under violations of exchangeability is also of interest and is discussed in the broader conformal literature.^23,29^ CP requires a non-conformity score that measures how atypical a candidate is for a given spectrum. For each spectrum *x* and candidate *c* ∈ 𝒜 (*x*), a non-conformity score is defined as *r*(*x, c*)∈ ℝ, where smaller values indicate higher conformity (i.e., a more plausible candidate) and larger values indicate lower conformity. The non-conformity scores used throughout the paper are constructed from the candidate scoring function *s*(*x, c*) introduced in Equation (1), and their specific forms are defined in Section 2.3. For each calibration pair (*x*_*i*_, *y*_*i*_), the calibration score is computed as *r*_*i*_ = *r*(*x*_*i*_, *y*_*i*_), where *y*_*i*_ denotes the true molecule associated with *x*_*i*_. Given an error level *α* ∈ (0, 1), split conformal prediction^30,31^ constructs a threshold *τ*_*α*_ as an empirical quantile of the calibration scores. Letting 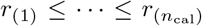 denote the sorted calibration scores, the standard split conformal quantile rule sets

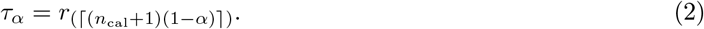

The prediction set for a test spectrum *x* is then obtained by including all candidates whose non-conformity score does not exceed this threshold,

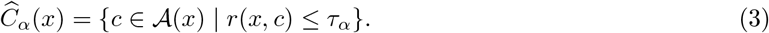

Different choices of *r*(*x, c*) can lead to different prediction set sizes, and can behave differently under heterogeneity in candidate-set size and score dispersion.^32,33^

The guarantee provided above controls the error rate on average over the test distribution and is a marginal guarantee,^12^ but it does not ensure that coverage is stable across subsets of spectra that differ in retrieval difficulty. To assess reliability across heterogeneous spectra, group-conditional coverage is considered with respect to a grouping function *g*: *𝒳*→ {1, …, *G*} that assigns each spectrum to one of *G* groups.^34^ The goal is for coverage to be close to the target within each group,

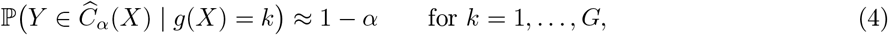

which provides a subgroup-level notion of reliability that is especially relevant in LC–MS/MS retrieval because candidate-set properties and score profiles can vary widely across spectra. This criterion is used throughout the paper to quantify coverage disparities across spectrum sub-populations and motivates the conditional conformal constructions introduced in Section 2.4. All conformal methods in this work are applied on top of a fixed, pre-trained retrieval model at confidence level 1 − *α* = 0.9.

### 2.3 Non-conformity scores

The CP methods in this work operate on the ranked candidate list defined in Section 2.1 and are therefore driven by the candidate scores *s*(*x, c*) introduced in Equation (1). When the retrieval model produces unnormalized scores *s*(*x, c*), these scores are converted into normalized values over the candidate set via a softmax transformation,

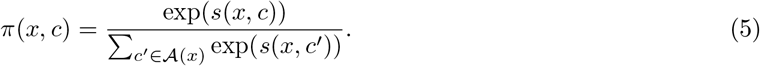

If the model already outputs probabilities over 𝒜 (*x*), then *π*(*x, c*) denotes those probabilities directly. In either case, larger *π*(*x, c*) indicates higher plausibility. A non-conformity score is a function *r*(*x, c*) ∈ ℝ such that smaller values correspond to more plausible candidates and larger values correspond to less plausible candidates. The simplest non-conformity score is the Least Ambiguous set-valued Classifier (LAC),^35^ defined as

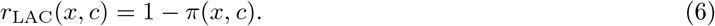

Under this score, a candidate is included in the prediction set whenever *π*(*x, c*) ≥ 1 − *τ*_*α*_. In addition to LAC, this work adopts Adaptive Prediction Sets (APS)^33^ and Regularized Adaptive Prediction Sets (RAPS),^32^ which extend the non-conformity score by accumulating plausibility along the ranked candidate list rather than thresholding each candidate independently. For each calibration pair (*x*_*i*_, *y*_*i*_) ∈ 𝒟_cal_, the score *r*_*i*_ = *r*(*x*_*i*_, *y*_*i*_) computed using any of the above definitions is used to determine conformal thresholds at a desired error level *α* via Equation (2).

In our retrieval setting, the normalized candidate scores *π*(*x, c*) used to compute these scores (Equation (5)) are derived from a similarity-based retrieval model rather than from a multi-class classifier. Consequently, the softmax probabilities are computed over the variable-length candidate set 𝒜 (*x*) for each spectrum, rather than over a fixed label set. Deterministic variants of APS and RAPS are used throughout for reproducibility.^32^ RAPS introduces hyperparameters (*k*_reg_, *λ*). We tune them on a held-out subset of the calibration data (size *n*_tune_ = 1000): *k*_reg_ is tuned based on the empirical ranks of the true candidates,^32^ and *λ* is chosen by grid search over {0.001, 0.01, 0.1, 0.2, 0.5} to minimize average set size while maintaining the target coverage on the tuning subset. The final conformal threshold 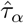 is then computed on the remaining calibration samples using Equation (2). For marginal conformal prediction, a single threshold is computed from the calibration set using the split conformal quantile rule (Equation (2)), and the prediction set for each test spectrum is constructed by thresholding *r*(*x, c*) as in Equation (3). For APS and RAPS, the resulting set corresponds to a top-ranked prefix of candidates because *r*(*x, c*_(*j*)_) is nondecreasing in rank *j*.

### 2.4 Conditional conformal prediction

Conditional conformal prediction is implemented through Mondrian (group-conditional) conformal prediction,^12,34,36^ with related extensions studying conditional guarantees under covariate shift,^21^ selection-conditional coverage,^37^ and localized conformal methods.^25^ A grouping function *g*(*x*) ∈ {1, …, *G*} partitions spectra into *G* covariate-defined sub-populations using only information available at prediction time, and a separate conformal threshold is estimated within each group. All conditional procedures rely on conditioning information available prior to observing the true label. We consider four conditioning variables: precursor mass, candidate set size |𝒜 (*x*) |, the maximum normalized score max_*c*∈𝒜(*x*)_ *π*(*x, c*), and a candidate-set similarity measure. Candidate-set similarity between spectra *x*_*i*_ and *x*_*j*_ is computed as the average pairwise Tanimoto similarity between binary fingerprints of candidates in their respective candidate sets:

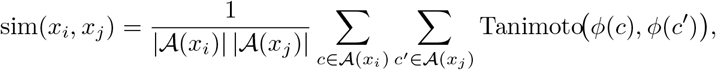

where *ϕ*(*c*) denotes the binary fingerprint of candidate *c*, and Tanimoto(·,·) is the standard Tanimoto coefficient.^38^ In addition to these four conditioning variables, we also consider the top-1/top-2 gap *π*(*x, c*_(1)_) − *π*(*x, c*_(2)_) as another candidate conditioning variable, because it provides a simple summary of how clearly the model separates its top prediction from the closest alternative. We then compute Pearson correlations among all five variables to assess overlap and avoid retaining redundant conditioning variables.

Let 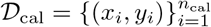 denote the calibration set and let *r*_*i*_ = *r*(*x*_*i*_, *y*_*i*_) be the scores computed using one of the non-conformity scores defined in Section 2.3. For each group *k* ∈ {1, …, *G*}, define the calibration index set

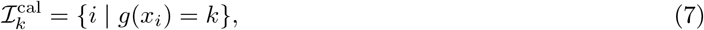

and the corresponding group calibration scores 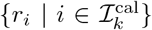. A group-specific threshold *τ*_*α,k*_ is obtained as the empirical (1 − *α*) quantile of the group calibration scores, using the split conformal quantile rule (Equation (2)) applied within the group. Under exchangeability of calibration and test samples within each group, this construction yields the group-conditional coverage guarantee in Equation (4) (up to the usual ≥ 1 − *α* inequality), as in Mondrian conformal prediction.^12,34^ For a test spectrum *x*, the prediction set is constructed using the threshold associated with its assigned group:

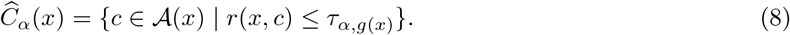

This framework reduces to marginal conformal prediction when *G* = 1 and illustrates a granularity–stability trade-off: increasing *G* yields more specific calibration but fewer calibration samples per group, which can make quantile estimates less stable.^13,24,29^ We consider two strategies for defining *g*(·) using the conditioning variables above.

#### Cluster-conditional conformal prediction (CCCP)

CCCP defines groups by clustering calibration spectra using a single conditioning variable at a time. Let *z*(*x*) ∈ ℝ denote the chosen conditioning variable. After standardization, a clustering algorithm is applied to 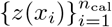 to obtain *G* clusters and thereby a grouping function *g*(*x*) ∈ {1, …, *G*}. Group-specific thresholds *τ*_*α,k*_ are computed from calibration scores within each cluster and applied to test spectra via Equation (8). In the experiments, we vary the number of clusters over *G* ∈ {10, 20, 30, 40, 50}.

#### Nearest-neighbor conformal prediction (CCP-NN)

CCP-NN replaces fixed clusters with a local calibration neighborhood around each test spectrum. Using the same choice of conditioning statistic *z*(*x*) as in CCCP, a distance is defined on *z*(*x*) and, for each test spectrum *x*, the indices of its *K* nearest calibration spectra are denoted by 𝒩_*K*_(*x*). A local threshold *τ*_*α*_(*x*) is computed as the empirical (1 − *α*) quantile of the local calibration scores {*r*_*i*_ |*i* ∈ 𝒩_*K*_(*x*)} using the split conformal quantile rule within the neighborhood, and the prediction set is obtained by replacing *τ*_*α,g*(*x*)_ in Equation (8) with *τ*_*α*_(*x*). Neighborhood size controls the locality–stability trade-off, and we vary it over *K* ∈ {100, 200, 300, 400, 500} in the experiments. Because CCP-NN defines a different neighborhood for each test spectrum, it does not induce shared, static groups. To report subgroup-level coverage and the conditional reliability metrics in Section2.5, test spectra are mapped to pseudo-groups by ordering and binning along the same conditioning statistic used to define neighborhoods. When neighborhoods are defined by candidate-set similarity, the ordering statistic is the average Tanimoto distance to the *K* nearest calibration neighbors. When neighborhoods are defined by precursor mass, candidate set size, or maximum softmax probability, the ordering statistic is the corresponding scalar value. Test spectra are sorted by this statistic and partitioned into consecutive blocks with sizes in {100, 200, 300, 400, 500} to match *K*, and group-wise summaries are computed within each block.

### 2.5 Experimental Setup

#### Dataset

Experiments use MassSpecGym,^1^ which contains 231,104 MS/MS spectra linked to 31,602 unique molecular structures. Each spectrum is represented by a feature vector *x* ∈ 𝒳 and is paired with its associated molecular structure *y* ∈ ℳ. MassSpecGym provides an official split of the data into training, validation, and test *folds*. These folds are designed to reduce structural leakage by grouping molecules using agglomerative clustering based on molecular-graph distances, and to balance key acquisition conditions across folds by stratifying spectra using instrument type, collision energy, ionization adduct, and molecule frequency. Candidate-based retrieval is evaluated using a predefined candidate set for each spectrum. For a spectrum *x*, let 𝒜 (*x*) ⊂ ℳ denote the candidate set, constructed by sampling molecules that match the precursor mass of the query spectrum up to a maximum of 256 candidates.^1^ As a result, candidate set sizes |𝒜 (*x*) | vary across spectra while remaining bounded by 256. The evaluation setting assumes that the true molecule belongs to the candidate set, i.e., *y* ∈ 𝒜 (*x*). Since the benchmark split does not include a dedicated calibration fold, an additional calibration split is created for conformal prediction by partitioning spectra according to the scenario definitions in the next paragraph. The calibration split is used only to estimate conformal thresholds, while the test split (or its scenario-specific derivative) is used only for final evaluation.

#### Scenarios and data splits

Evaluation is performed under three scenarios that control how spectra are assigned to the training, validation, calibration, and test splits (Table 1). Let 𝒟 denote the full dataset and let 𝒟_tr_, 𝒟_val_, 𝒟_cal_, and 𝒟_test_ denote the training, validation, calibration, and test splits, respectively. The retrieval model is fit using 𝒟_tr_ and selected using 𝒟_val_. Conformal thresholds are computed only from 𝒟_cal_, and all reported results are computed only on 𝒟_test_. In the defined scenarios, MassSpecGym folds are used either directly or as sources from which additional splits are created. All scenarios are defined at the spectrum level, but when a distribution shift is required, the split is performed at the level of molecular clusters (so that spectra associated with molecules from different clusters fall into different splits). When matched distributions are required, spectra are split randomly within the relevant pool. Scenario 1 (S1) is an IID reference setting in which all four splits are obtained by randomly partitioning the full dataset. The training split contains 73.6% of samples, matching the size of the provisional training partition from the initial cluster-based split used in the MassSpecGym construction pipeline. Unlike the final MassSpecGym benchmark, we do not apply the additional post-processing step that reassigns random validation and test clusters to the training fold, because this would not preserve the scenario-specific distributional relationships defined in our study. The remaining spectra are randomly divided into 𝒟_val_, 𝒟_cal_, and 𝒟_test_. Scenario 2 (S2) introduces a distribution shift for model fitting while keeping calibration and testing aligned. Training and validation are drawn from different molecular clusters following the MassSpecGym cluster structure, and 𝒟_cal_ and 𝒟_test_ are obtained by randomly splitting the original MassSpecGym test fold. Scenario 3 (S3) introduces the strongest mismatch by additionally breaking the alignment between calibration and test data. Training and validation are identical to S2, but calibration and test samples are drawn from different molecular clusters. In this scenario, exchangeability between 𝒟_cal_ and 𝒟_test_ is not expected to hold, and deviations from the nominal conformal coverage level are anticipated.^23,29^

**Table 1:**
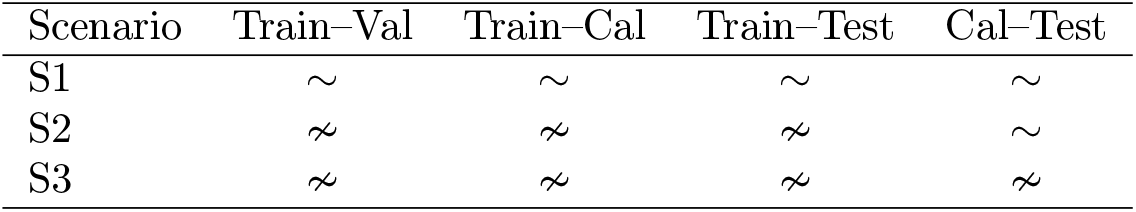
Scenario definitions. Each cell indicates whether the two splits share the same distribution (∼) or are drawn from different molecular clusters (≁).

#### Evaluation metrics

Evaluation is performed on the test split 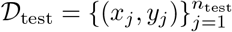, where *n*_test_ = |𝒟_test_| denotes the number of test spectra. The main outcomes are coverage and efficiency of the prediction sets 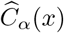 constructed by marginal or conditional conformal prediction. Empirical marginal coverage is computed as

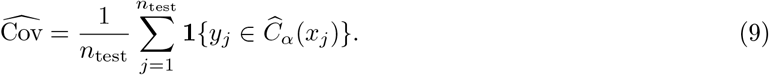

For conditional conformal prediction, group-wise coverage is computed with respect to the grouping function *g*(*x*) ∈ {1, …, *G*} introduced in Section 2.4, where *G* denotes the total number of groups. For each group *k* ∈ {1, …, *G*}, let I_*k*_ = {*j* | *g*(*x*_*j*_) = *k*} and *n*_*k*_ = |*ℐ*_*k*_|, and define

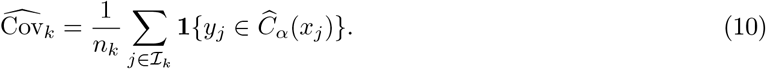

To summarize coverage disparities across groups, the mean absolute coverage gap (MACG)^24^ is reported,

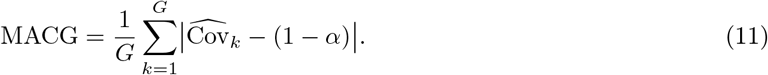

where smaller values indicate more uniform group-wise coverage around the target level 1 − *α*. Efficiency is summarized by the average prediction set size,

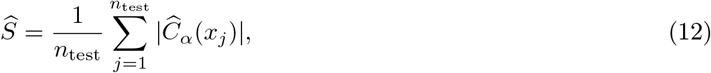

and by the mean relative size that accounts for varying candidate set sizes,

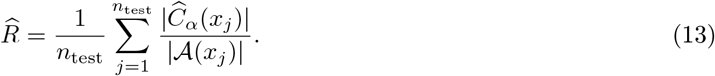

When relevant, group-wise averages of 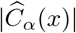 and 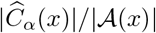 are also reported to assess how efficiency varies across spectrum sub-populations. Figure 1 provides a schematic overview of the complete framework.

**Figure 1.**
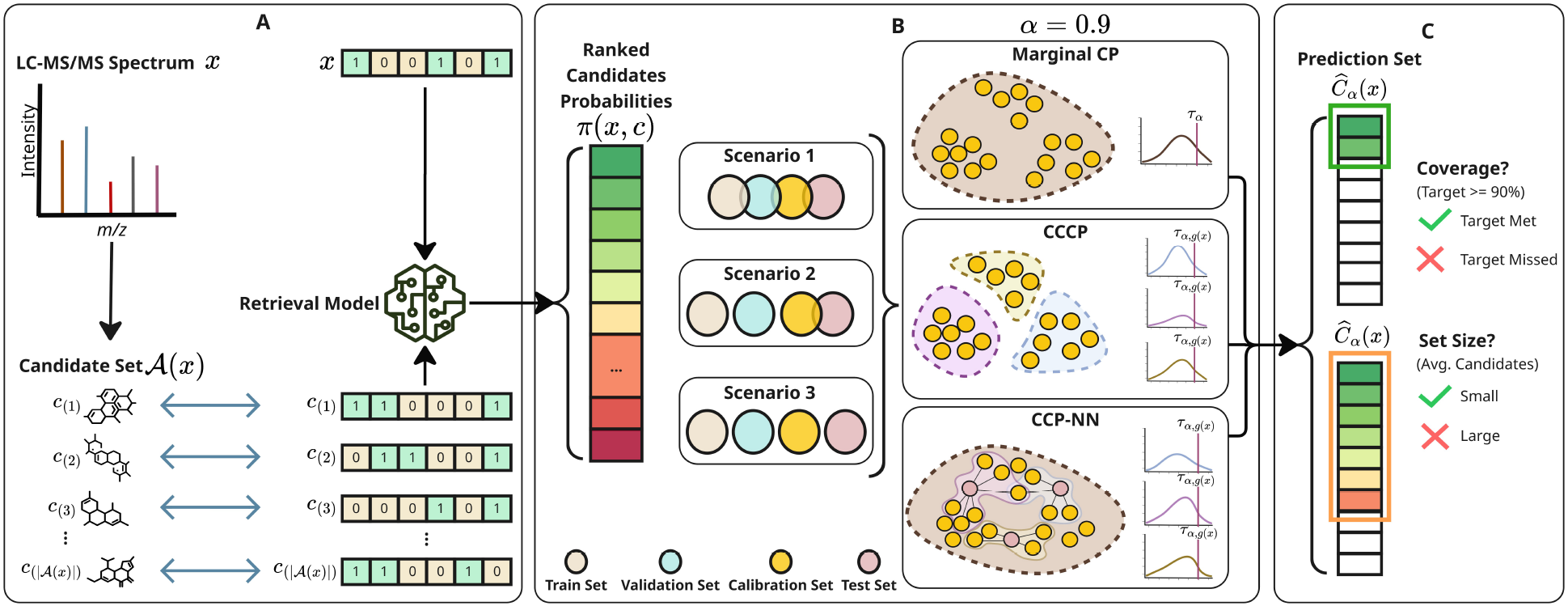
Overview of the uncertainty-aware molecular retrieval framework. (A) An input spectrum *x* is mapped to a predicted fingerprint *f* (*x*) by the retrieval model and compared to candidate fingerprints *ϕ*(*c*) in the mass-filtered set 𝒜 (*x*). Candidates are scored via *s*(*x, c*) = sim(*f* (*x*), *ϕ*(*c*)) and transformed into normalized probabilities *π*(*x, c*) via softmax, yielding a ranked list. (B) Non-conformity scores (LAC, APS, RAPS) are computed from *π*(*x, c*) on the calibration set 𝒟_cal_ and converted into prediction sets at target coverage 1 − *α* = 0.9. Marginal conformal prediction uses a single global threshold *τ*_*α*_, whereas cluster-conditional conformal prediction (CCCP) estimates group-specific thresholds *τ*_*α,k*_ via a grouping function *g*(*x*). Meanwhile, nearest-neighbor conformal prediction (CCP-NN) computes local thresholds *τ*_*α*_(*x*) from the *K* nearest calibration spectra 𝒩_*K*_(*x*). The procedure is evaluated under three scenario definitions that vary the alignment between training, calibration, and test data. (C) For each test spectrum, conformal calibration yields a prediction 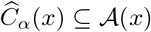 that retains only candidates with *r*(*x, c*) ≤ *τ*. Performance is assessed via empirical coverage 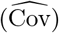, mean relative set size 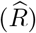, and mean absolute coverage gap (MACG).

## 3 Results and discussion

We organize the results around three questions. First, we examine how retrieval difficulty varies across the three scenarios (Section 3.1), which contextualizes the conformal prediction analysis that follows. Second we assess how well marginal conformal prediction achieves the target coverage and at what efficiency cost for LAC, APS, and RAPS (Section 3.2). Third, we investigate whether conditional conformal prediction can reduce subgroup coverage disparities, including how conditioning variables behave (Section 3.3) and how CCCP and CCP-NN trade reliability for set size under distribution shift (Section 3.4).

### 3.1 Retrieval baseline and scenario difficulty

Table 2 summarizes baseline retrieval performance across scenarios. We report top-*k* accuracy from the ranking induced by *s*(*x, c*) (Equation (1) in Section 2.1), together with indicators of candidate-set size and score concentration. In Scenario 1, top-1 accuracy reaches 87.1% and the average maximum softmax probability is 0.72, indicating that the model often assigns a dominant score to the correct candidate. In the more demanding shifted settings (Scenarios 2 and 3), top-1 accuracy drops sharply to approximately 10% and the average maximum softmax probability falls to roughly 0.3. The average softmax gap between the top two candidates decreases from 0.67 in S1 to approximately 0.19 in S2 and S3, which confirms that the score distributions become substantially flatter and more ambiguous under shift. Consistently, the average rank of the true molecule increases from 3.7 in S1 to 61.7 in S2 and 63.9 in S3, indicating that correct molecules are typically buried much deeper in the ranking. Candidate sets are large in all scenarios, with an average of 212 candidates per spectrum in S1 and 168 in S2 and S3. S2 and S3 exhibit similar baseline performance because they share the same trained model, while the distinction between them lies in whether calibration and test are aligned (S2) or deliberately mismatched in S3. Overall, these statistics quantify a strong asymmetry across scenarios that is essential for interpreting the conformal prediction results, since the difficulty of the underlying retrieval task directly determines how informative the non-conformity scores are about the rank position of the true molecule.

**Table 2:** Retrieval baseline results across scenarios. Top-*k* accuracy, average rank of the true molecule, average candidate set size, average maximum softmax probability, and average softmax gap between the top two candidates.

| Scenario | Top-1 | Top-5 | Top-20 | Avg Rank | Avg Cands | Avg Max Softmax | Avg Softmax Gap |
| --- | --- | --- | --- | --- | --- | --- | --- |
| S1 | 0.871 | 0.929 | 0.964 | 3.66 | 211.59 | 0.724 | 0.673 |
| S2 | 0.100 | 0.204 | 0.403 | 61.67 | 168.17 | 0.306 | 0.188 |
| S3 | 0.101 | 0.202 | 0.386 | 63.92 | 167.54 | 0.307 | 0.194 |

It is also worth noting that these scenarios are controlled extremes rather than a direct mirror of typical metabolomics data. In any realistic LC–MS/MS experiment, molecules overlapping with the training dataset may be present (i.e., as expected in S1), while others may be OOD, such as in S2 and S3. MassSpecGym is a curated benchmark and it cannot cover all practical shifts, but its structure-disjoint splits make it useful for controlled stress tests.^1^ The conformal prediction results can therefore be read as bracketing expected behavior, with performance often falling between an easier setting (S1) and a hard, mismatched setting (S3), depending on how different the molecules and conditions are from the training data. The retrieval model used here is a fingerprint-based MLP trained with a listwise ranking objective,^28^ and it was competitive at the time of this study. The field is moving quickly and MassSpecGym has helped drive new retrieval architectures.^6^ Recent joint embedding methods such as JESTR^10^ and MVP^39^ report slightly higher retrieval accuracy on the same benchmark, so the gap between S1 and S2 or S3 may shrink with stronger models. Even so, the conformal framework studied here is model-agnostic. It uses only the scored candidate list and does not need model internals or retraining. If retrieval scores improve, prediction sets should become smaller and more useful, so progress in architectures is complementary to conformal calibration. The results that follow should thus be interpreted in two ways, as a characterization of conformal prediction behavior for the current retrieval model, and as a conservative baseline for the efficiency gains achievable when the same conformal methods are applied to stronger future models.

### 3.2 Marginal conformal prediction across scenarios

We evaluate marginal conformal prediction at target coverage 1 − *α* = 0.9 using the three non-conformity scores defined in Section 2.3. These scores are representative of common design choices in split conformal prediction and are particularly relevant when candidate sets are large, since the score definition determines the shape of the resulting prediction sets and can substantially affect set sizes under diffuse plausibility distributions.^40–42^ LAC directly thresholds candidate plausibility, APS aggregates plausibility along the ranked list, and RAPS regularizes this cumulative construction to reduce sensitivity to low-ranked candidates that may be dominated by noise.^32^ Alternative constructions exist, including rank-based scores that discard probability values entirely^43^ and SAPS, which retains only the maximum softmax probability to reduce sensitivity to miscalibrated tails.^44^ Table 3 summarizes empirical marginal coverage and efficiency for LAC, APS, and RAPS across all three scenarios. In S1, where calibration and test data are exchangeable, all three scores achieve marginal coverage at or above the nominal 90% level, confirming the finite-sample guarantee. All three scores produce very small prediction sets in this setting: APS retains an average of 1.5 candidates (1.7% of the candidate set), LAC retains 1.6 candidates (2.1%), and RAPS retains 3.1 candidates (3.5%). The near-identical efficiency of LAC and APS reflects the high model confidence in S1: when the softmax distribution is strongly peaked on a single candidate, both the threshold-based LAC score and the cumulative APS score include essentially only the top-ranked candidate. RAPS produces slightly larger sets and exhibits mild overcoverage (91.7% instead of the target 90%), which is a known consequence of the discrete nature of cumulative scores combined with its regularization penalty: when the model is confident, the cumulative mass jumps sharply at low ranks and the penalty discourages splitting this jump, making it difficult to fine-tune the threshold to achieve exact 1 − *α* coverage. In S2 and S3, all prediction sets expand dramatically: relative sizes exceed 80% of the candidate set for all three scores, compared to 2–4% in S1. This expansion is expected because the retrieval model generalizes poorly to novel molecular clusters, and the softmax distributions become flatter and less informative. When scores are diffuse, the conformal threshold must be set high enough to capture the true molecule in 90% of cases, which unavoidably includes most of the candidate set. In S2, where calibration and test data remain exchangeable, all three scores maintain valid marginal coverage (0.898–0.902). In S3, which breaks exchangeability between calibration and test, coverage drops below the nominal level for all scores, most notably for APS (0.881) and LAC (0.884), illustrating that the finite-sample guarantee does not hold when calibration and test data come from different molecular-cluster distributions. RAPS proves more robust under this shift, maintaining coverage closest to the target at 0.895, likely because its regularization produces inherently more conservative thresholds. The efficiency differences across scores observed in S1 largely collapse in S2 and S3, where all three scores produce relative set sizes in the 80–83% range. RAPS retains a slight advantage in absolute set size (125.8 in S2, 127.0 in S3) compared to LAC (132.9, 130.5) and APS (134.9, 132.8), but the practical difference is small given that all methods must retain the majority of candidates. This convergence reflects a fundamental limitation: when the retrieval model cannot reliably separate the true molecule from its competitors, the softmax distributions become too flat for any non-conformity score to distinguish correct from incorrect candidates, and the conformal threshold must admit most of the candidate set to achieve coverage.

**Table 3:** Marginal conformal prediction at 1 − *α* = 0.9: empirical coverage, average prediction set size, and mean relative set size 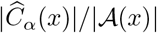 across scenarios.

| Scenario | Score | Coverage | Avg Set Size | Rel. Size (%) |
| --- | --- | --- | --- | --- |
| S1 | LAC | 0.901 | 1.6 | 2.1 |
|  | APS | 0.900 | 1.5 | 1.7 |
|  | RAPS | 0.917 | 3.1 | 3.5 |
| S2 | LAC | 0.899 | 132.9 | 82.3 |
|  | APS | 0.898 | 134.9 | 81.1 |
|  | RAPS | 0.902 | 125.8 | 83.2 |
| S3 | LAC | 0.884 | 130.5 | 81.8 |
|  | APS | 0.881 | 132.8 | 80.6 |
|  | RAPS | 0.895 | 127.0 | 82.3 |

### 3.3 Conditioning variables and correlation analysis

Before presenting conditional CP results, we examine which conditioning variables (defined in Section 2.4) provide complementary information about retrieval difficulty and how redundancy between them is handled.

Figure 2 summarizes this analysis for Scenario 1 using the pooled calibration and test splits, and Figure S1 reports the same Pearson correlation heatmaps for all scenarios and splits. Figure 2(a) shows that maximum softmax probability and the top-1/top-2 softmax gap are almost identical (*ρ* = 0.98), so we keep max softmax as the confidence variable and drop the softmax gap. Candidate set size and precursor mass are negatively correlated (*ρ* = − 0.50), so heavier molecules usually come with smaller candidate sets. This pattern is consistent with how chemical databases are populated because low-mass compounds are far more common, which means low-mass query spectra often match many same-mass molecules and produce large, structurally diverse candidate sets, whereas high-mass spectra have fewer same-mass matches and therefore yield smaller sets. This difference directly shapes retrieval difficulty, since larger and more diverse candidate sets make it harder to rank the correct structure near the top. In contrast, candidate set size is almost uncorrelated with max softmax (*ρ* ≈ − 0.02), suggesting that candidate set size and model confidence capture different aspects of difficulty. Max softmax is mildly negatively correlated with precursor mass (*ρ* = − 0.26), indicating that heavier precursors are associated with flatter, less decisive score distributions.

**Figure 2.**
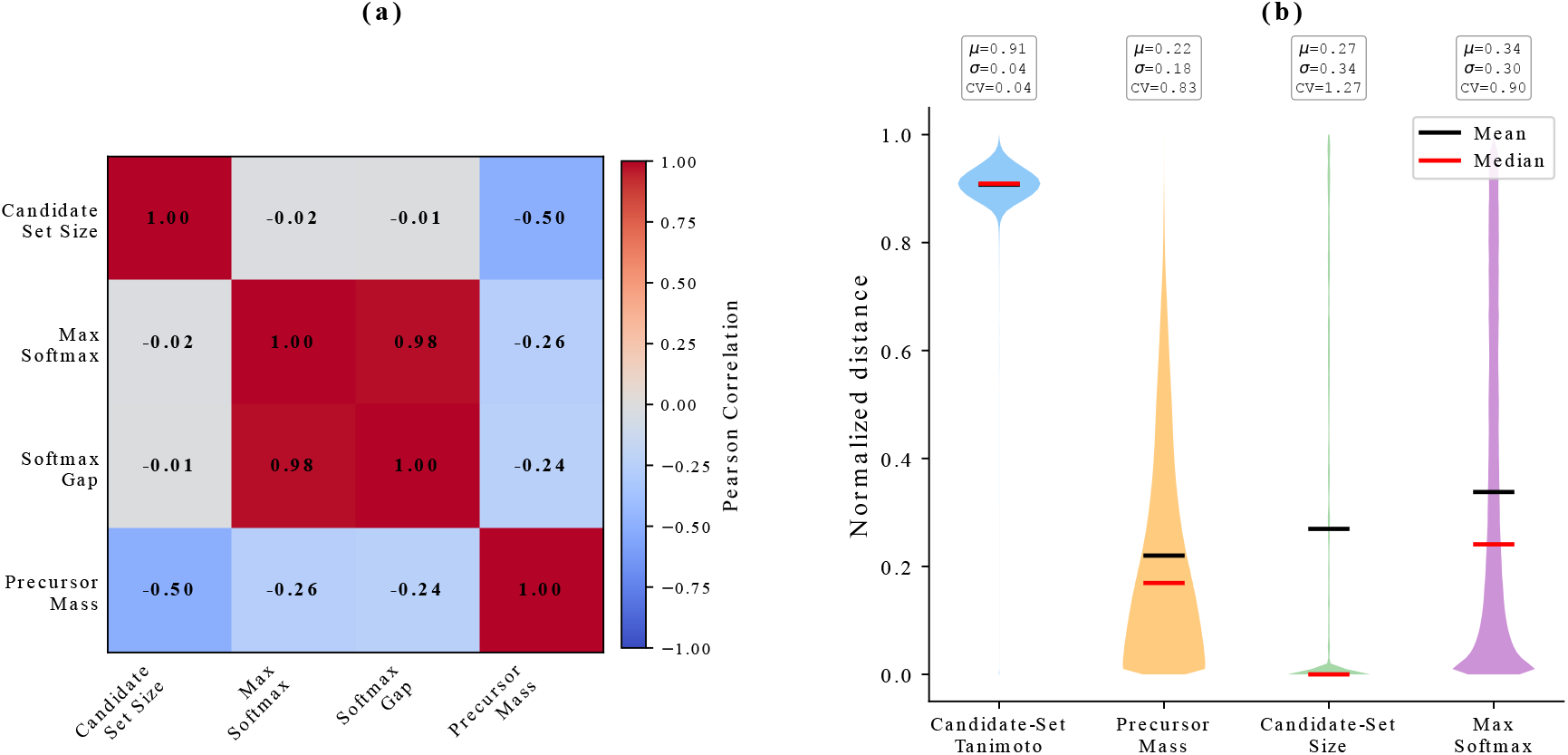
Conditioning-variable diagnostics for Scenario 1 (calib+test pooled). (a) Pearson correlations among scalar conditioning variables (candidate set size, max softmax, softmax gap, precursor mass). (b) Violin plots of min–max normalized pairwise distances for the retained conditioning variables, with mean (*µ*), standard deviation (*σ*), and coefficient of variation (CV) reported above each violin. Horizontal black and red bars indicate mean and median, respectively.

Figure 2(b) compares the geometry induced by each retained conditioning variable after min–max normalization. Candidate-set dissimilarity is tightly concentrated near high distances (*µ* = 0.91, *σ* = 0.04), which limits its ability to form informative neighborhoods or balanced clusters. Precursor mass distances are more dispersed (*µ* = 0.22, *σ* = 0.18), while candidate set size and max softmax show broader and more variable distance distributions (*µ* = 0.27, *σ* = 0.34 and *µ* = 0.34, *σ* = 0.30), supporting their use for separating spectra by difficulty. Figure S2 shows the corresponding distance distributions before normalization, reported in their original units.

The cluster quality analysis in Figure 3 assesses how the retained conditioning variables behave when used for agglomerative clustering at *G* ∈ {10, 20, 30, 40, 50}. Candidate-set dissimilarity again stands out as problematic, producing highly unbalanced partitions with many clusters falling below the 100-sample threshold as *G* increases. By contrast, precursor mass and max softmax yield substantially more even cluster sizes, while candidate set size degrades as granularity increases. These patterns anticipate the CCCP results in Section 3.4: conditioning variables that generate many undersized clusters lead to unstable within-cluster quantile estimates, which in turn inflates MACG and degrades subgroup reliability.

**Figure 3.**
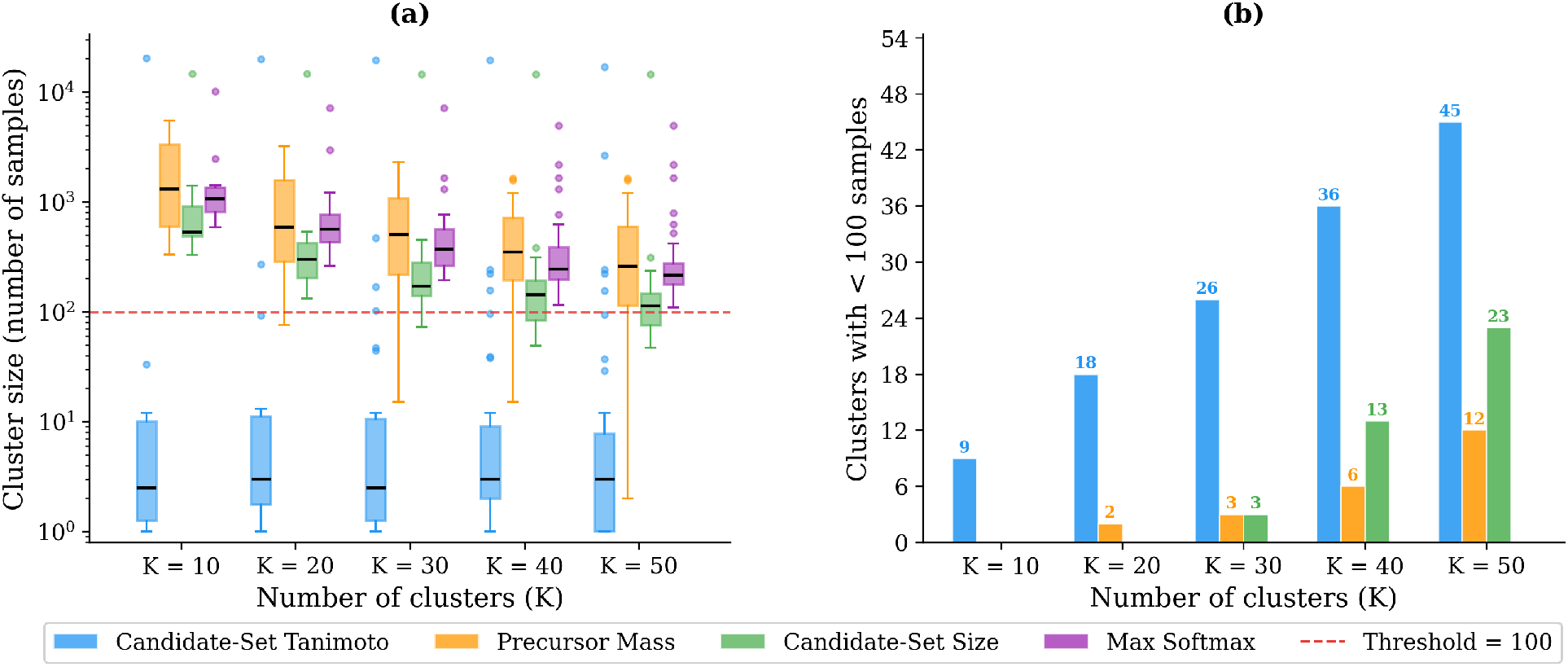
Cluster quality across four conditioning variables. (a) Distribution of cluster sizes for *G* ∈ {10, 20, 30, 40, 50}. The dashed red line marks the 100-sample threshold. (b) Number of clusters with fewer than 100 calibration samples.

### 3.4 Conditional conformal prediction: CCCP vs. CCP-NN

We evaluate conditional conformal prediction through both CCCP and CCP-NN at target coverage 1 − *α* = 0.9, using all three non-conformity scores and the four retained conditioning variables. Uniformity of coverage across sub-populations is summarized by MACG (Equation (11)), and efficiency is assessed by the mean relative prediction set size 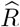 (Equation (13)). Figures 4 and 5 report results for CCCP with *G* = 10 clusters and CCP-NN with neighborhood size *K* = 100. Complete results across *G* ∈ {10, 20, 30, 40, 50} and *K* ∈ {100, 200, 300, 400, 500} are reported in Supplementary Figures S3 and S4. Also, mean conditional coverage values, computed as 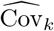 in Equation (10), are reported for all settings in Table S1.

**Figure 4.**
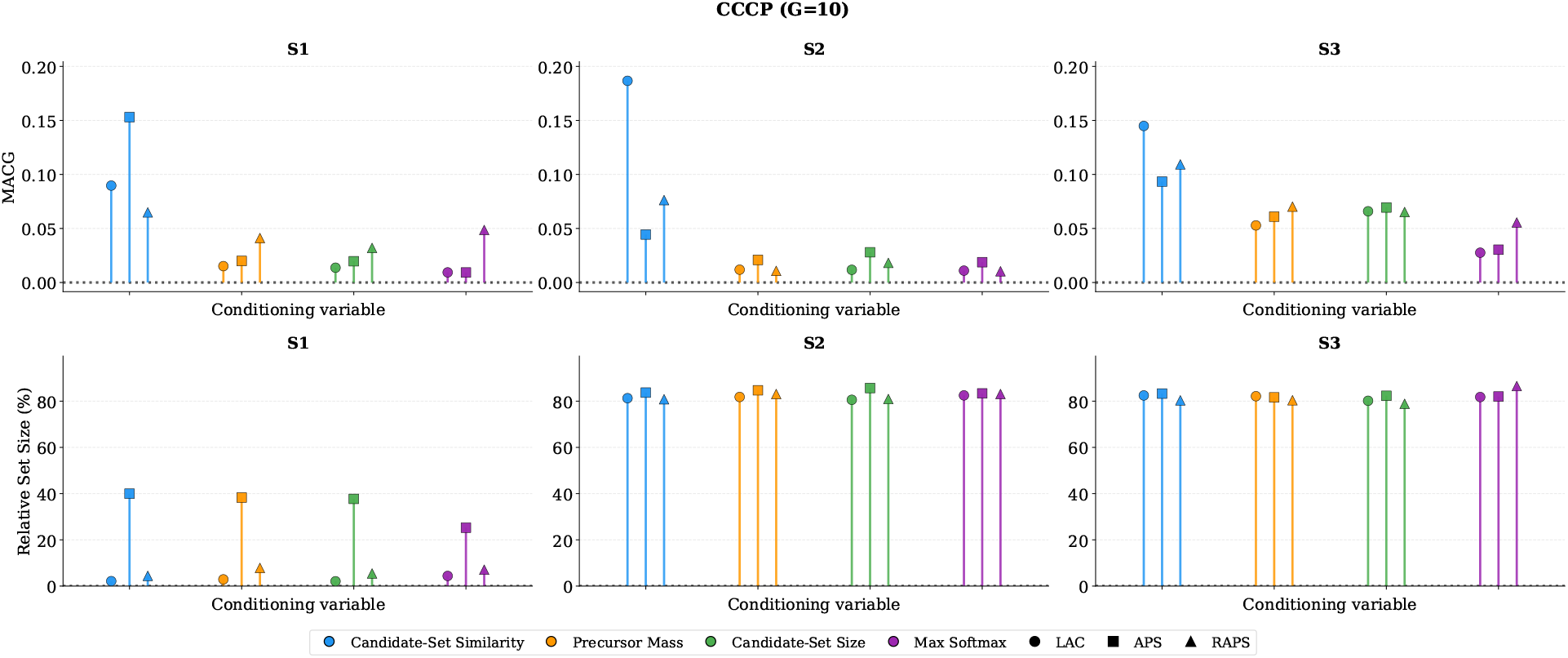
CCCP at 1 − *α* = 0.9 with *G* = 10. Top: MACG. Colors denote conditioning variables and markers denote scores (LAC/APS/RAPS). Bottom: mean relative set size 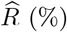 (%).

**Figure 5.**
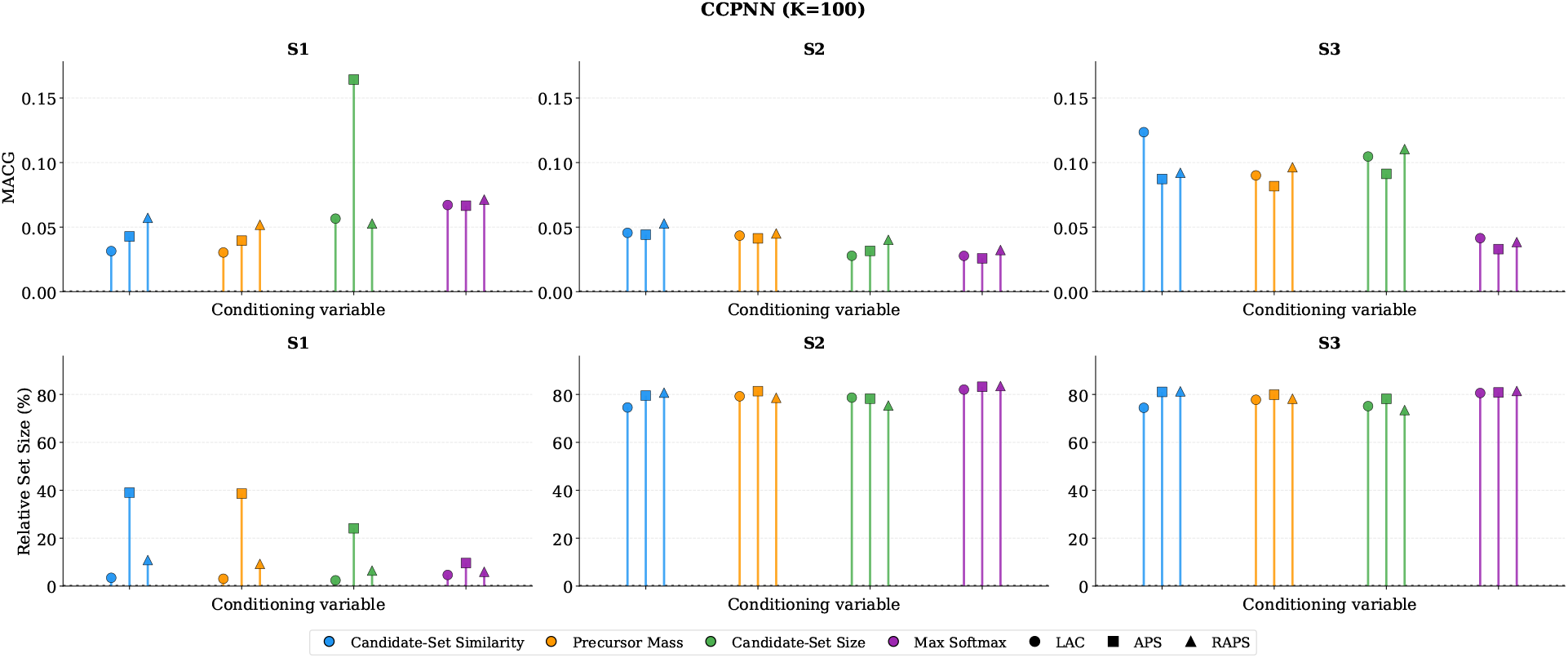
CCP-NN at 1 − *α* = 0.9 with *K* = 100. Top: MACG. Colors denote conditioning variables and markers denote scores (LAC/APS/RAPS). Bottom: mean relative set size 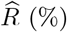 (%).

The choice of conditioning variable has the largest and most consistent effect on conditional coverage quality. Almost across all configurations, max softmax achieves the lowest MACG, making it the most informative variable for reducing subgroup coverage disparities. In S1, conditioning on max softmax with CCCP (*G* = 10) yields MACG values of 0.009 for both LAC and APS. Even under the distribution shift of S3, max softmax retains a clear advantage, achieving MACG values of 0.028 (LAC), 0.030 (APS), and 0.056 (RAPS) with CCCP at *G* = 10, and 0.035 (LAC), 0.024 (APS), and 0.030 (RAPS) with CCP-NN at the best *K*. In both methods, these values are roughly half the MACG obtained by the next-best variable. This result is consistent with the cluster quality analysis in Section 3.3: max softmax produces the most balanced cluster partitions (Figure 3) and has broad distance spread (Figure 2), both of which support stable within-group quantile estimation. More fundamentally, max softmax directly reflects how confident the model is about each spectrum, which makes it the most natural proxy for retrieval difficulty. By calibrating thresholds separately within sub-populations defined by this variable, the conformal procedure adapts to distinct difficulty patterns. Precursor mass and candidate set size achieve intermediate MACG values. With CCCP (*G* = 10), precursor mass yields MACG of 0.015 (LAC) and 0.020 (APS) in S1, and 0.012 (LAC) and 0.021 (APS) in S2, degrading to 0.053–0.070 in S3. Candidate set size shows a similar pattern, with MACG values of 0.014–0.032 in S1, 0.012–0.028 in S2, and 0.065–0.069 in S3. These variables capture different aspects of retrieval difficulty—mass constrains the chemical search space, and candidate set size determines the combinatorial challenge—but they are less directly connected to the model’s scoring behavior than max softmax. The candidate-set similarity variable performs most poorly, consistently producing the highest MACG. In S1, CCCP with candidate-set similarity and APS yields a MACG of 0.153, an order of magnitude worse than the 0.009 achieved by max softmax. Under S3 the gap persists, with MACG reaching 0.093 for APS compared to 0.030 with max softmax. This poor performance follows directly from the unbalanced cluster structure in Figure 3: the narrow, high-distance concentration of Tanimoto distances causes many small clusters with insufficient calibration samples for reliable quantile estimation. In CCP-NN, candidate-set similarity performs somewhat better because the nearest-neighbor construction does not require well-separated clusters, but it still falls behind mass and max softmax in most configurations, except in S1.

Turning to the non-conformity scores, conditional calibration reveals important differences from the marginal case. A notable observation is that conditional CP in S1 produces larger prediction sets than marginal CP, despite the IID setting. While marginal CP achieves sets of 1.5–3.1 candidates (1.7–3.5%, Table 3), the best CCCP configurations with max softmax (*G* = 10) yield 6.2 candidates (4.4%) for LAC, 54.5 candidates (25.3%) for APS, and 9.1 candidates (7.1%) for RAPS. This expansion occurs because group-wise calibration must set thresholds that ensure coverage within each sub-population separately: groups with lower model confidence need higher thresholds, and these inflate prediction sets even for spectra that would have been easy under global calibration. The expansion is most pronounced for APS, whose cumulative score accumulates many low-ranked candidates once the threshold is raised. Among the scores in S1, LAC achieves the best balance of reliability and efficiency: MACG of 0.009 with a relative set size of 4.4%. APS matches the MACG (0.009) but at much larger set sizes (25.3%). RAPS has a higher MACG (0.049) with max softmax conditioning in S1, because its overcoverage tendency (91.7% marginal coverage) interacts with group-wise calibration to produce systematically overcovered sub-populations. The best RAPS configuration in S1 instead uses candidate set size conditioning, with a MACG of 0.032. In S2 and S3, efficiency differences across scores largely disappear, with all relative set sizes in the 80–87% range, but meaningful MACG differences remain. In S2, RAPS achieves the lowest MACG with CCCP max softmax (0.010), followed by LAC (0.011) and APS (0.019). In S3, APS achieves the lowest MACG with CCP-NN max softmax (0.024), followed by RAPS (0.030) and LAC (0.035). These shifts in ranking between scenarios show that no single score dominates everywhere. Instead, the conditioning variable and conditioning algorithm choice have a larger and more consistent effect on MACG than the score choice.

Comparing the two algorithms directly, using max softmax at the best hyperparameter for each, reveals a clear scenario-dependent pattern. In S1, CCCP achieves much lower MACG than CCP-NN across all scores. For APS, CCCP yields MACG = 0.009 at *G* = 10, whereas CCP-NN gives 0.063 at *K* = 400, which corresponds to roughly a sevenfold improvement. Similarly, for LAC the gap is 0.009 versus 0.065, and for RAPS it is 0.048 versus 0.069. CCCP works well here because it constructs fixed clusters and computes a separate quantile within each, which is effective when calibration and test data are well-aligned. CCP-NN, by contrast, produces systematic overcoverage in S1 (mean coverage around 96%), inflating the MACG by deviating upward from the 90% target. In S2, the gap narrows. CCCP retains the advantage for LAC (0.011 vs. 0.016) and RAPS (0.010 vs. 0.022), but CCP-NN achieves a slightly lower MACG for APS (0.017 vs. 0.019). In S3, where exchangeability is broken, CCP-NN outperforms CCCP for APS (0.024 vs. 0.030) and RAPS (0.030 vs. 0.036), though CCCP retains its advantage for LAC (0.028 vs. 0.035). The advantage of CCP-NN under shift is intuitive: fixed clusters formed on calibration data may not match the difficulty structure of test spectra, while CCP-NN selects the most similar calibration samples for each test point, partially compensating for distributional mismatch. Overall, CCCP should be preferred when calibration and test data are well-aligned, while CCP-NN offers a useful alternative under distribution shift, particularly for APS and RAPS.

The number of clusters *G* and neighborhood size *K* control the granularity–stability trade-off. For APS with max softmax, increasing *G* from 10 to 50 in CCCP monotonically increases MACG in all scenarios: from 0.009 to 0.024 in S1, from 0.019 to 0.031 in S2, and from 0.030 to 0.043 in S3. Fewer clusters mean more calibration samples per group, leading to more stable quantile estimates. The efficiency impact is modest: in S1, relative set sizes decrease slightly from 25.3% to 20.5% as *G* grows, but this small gain is offset by the degraded coverage reliability. In S2 and S3, set sizes stay nearly constant across *G* (83–84% and 80–82%), confirming that cluster granularity mainly affects coverage uniformity rather than efficiency under shift. These results suggest *G* = 10 as the best overall choice when the calibration set contains approximately 20,000 spectra. For CCP-NN, MACG is less sensitive to *K* and the patterns depend on the scenario. In S1, MACG remains flat across *K* (0.063–0.067) because the overcoverage effect dominates. In S2, MACG drops overall from 0.026 (*K* = 100) to 0.017 (*K* = 500), though the decline is not strictly monotonic, indicating that larger neighborhoods generally stabilize the local quantile estimates. In S3, the pattern is non-monotonic: MACG drops from 0.033 (*K* = 100) to 0.024 (*K* = 300) and then rises slightly to 0.027 (*K* = 500), reflecting a tension between locality and stability—small neighborhoods adapt closely to the test point but yield noisy thresholds, while large neighborhoods smooth out the shift too aggressively.

Finally, the reliability–efficiency trade-off behaves qualitatively differently across scenarios, driven by the quality of the retrieval model. In S1, conditional CP achieves high reliability (MACG as low as 0.009) but at a cost relative to marginal CP: prediction sets expand from 1.5–3.1 candidates to 6.2–54.5. Despite this increase, sets remain small in absolute terms (at most 25% for APS, below 8% for LAC and RAPS), so the trade-off is favorable. In S2, conditional CP still reduces MACG effectively (0.010–0.019 for the best configurations), and the efficiency picture changes: all methods produce sets covering 81–84% of candidates, comparable to marginal CP (81–83%, Table 3). This means conditional calibration improves coverage uniformity at essentially no additional efficiency cost, because sets are already near-maximal. In S3, MACG increases further (0.024–0.030), and marginal coverage drops below 90% for all scores (Table 3), confirming that the conformal guarantee breaks under violated exchangeability. CCP-NN is preferable in this scenario for APS and RAPS because its local neighborhoods can partially compensate for the mismatch between calibration and test data.

### 3.5 Summary of key findings

The experiments show fairly clear trends. First, the maximum softmax probability is the most useful conditioning variable. It gives the lowest MACG in almost all settings, for both CCCP and CCP-NN, because it directly reflects model confidence and separates easy vs. hard spectra better than precursor mass, candidate-set size, or candidate-set similarity. In contrast, candidate-set similarity performs poorly for CCCP because it creates many very small clusters, which makes group-wise quantiles unstable. Second, the non-conformity score mainly matters when the retrieval model is strong. In S1, LAC is much more efficient than APS (prediction sets about 6–10 × smaller), while RAPS is slightly more conservative. Under distribution shift (S2 and S3), these efficiency differences largely disappear: all scores produce very large sets (typically *>* 70% of the candidate pool), because the ranking is ambiguous and the softmax distributions are flat. Third, CCCP and CCP-NN have complementary strengths. CCCP gives more stable conditional coverage when calibration and test are aligned, as in S1 and S2, whereas CCP-NN is more robust when calibration and test are mismatched, as in S3, because local neighborhoods can better adapt to the test distribution. Also, increasing conditioning granularity is not always beneficial. For CCCP, *G* = 10 is consistently the best choice because it keeps enough calibration samples per group. For CCP-NN, the best neighborhood size depends on the scenario, and intermediate values, such as *K* = 200–400, often work best under shift. Overall, these results show that conformal prediction improves reliability, especially subgroup stability, while its ability to reduce candidates depends on how informative the retrieval scores are. In S1, marginal CP reduces the candidate set to about 2% on average while maintaining 90% coverage. In the more challenging shifted settings, namely S2 and S3, prediction sets are necessarily larger, but they still remove a meaningful fraction of candidates, typically up to 25%, without sacrificing the target coverage.

## 4 Conclusion

Reliably identifying small molecules from LC–MS/MS data is central to metabolomics workflows ranging from biomarker discovery and drug metabolism studies to environmental monitoring and natural product characterization.^2,3^ Standard retrieval pipelines return a ranked candidate list for each spectrum but provide no per-spectrum reliability statement. In this work, we showed that conformal prediction can bridge this gap by converting any scored candidate ranking into a prediction set with a user-chosen error rate. When calibration and test data are well-aligned, these sets are small, often containing only a few candidates, and they give practitioners a concise and trustworthy shortlist. Conditional calibration using clustering-based and nearest-neighbor approaches can further improve stability across spectra of varying difficulty, reducing the risk that particular subgroups are consistently over- or under-covered. In practical terms, this enables reporting, for each spectrum, a set of plausible molecular structures accompanied by an explicit and interpretable confidence level, rather than a single best guess with no uncertainty quantification.

A key feature of this framework is that it is architecture-agnostic because conformal prediction operates on the output scores of the retrieval model and requires no modification to the model itself. This is particularly relevant as computational mass spectrometry is evolving rapidly, from fingerprint-based approaches such as CSI:FingerID^9^ and formula-transformer models such as MIST,^11^ to joint embedding methods that map spectra and molecules into a shared latent space, including JESTR,^10^ GMLR,^45^ and SpecBridge.^46^ Because the conformal prediction only requires a scored candidate list, it can be applied on top of these systems without retraining. The same principle extends beyond LC–MS/MS retrieval to any identification task that produces ranked hypotheses from mass spectral data, including MALDI-TOF-based microbial identification,^19^ GC–MS workflows, and *de novo* generation pipelines^8,47^ in which generated candidates are subsequently re-ranked.

Several limitations should be acknowledged. First, conformal prediction cannot recover efficiency when the retrieval scores themselves provide weak separation between candidates, so under distribution shift prediction sets grow because the ranking itself becomes ambiguous. Second, while MassSpecGym provides a carefully curated benchmark with structure-disjoint splits that enable controlled stress tests,^1^ it cannot capture the full diversity of practical shifts, such as instrument drift, novel chemistries, sample-matrix effects, and evolving databases. Third, our results are obtained under the capped candidate regime considered by MassSpecGym, with up to 256 candidates, and assume the true molecule is in the candidate pool, so larger candidate sets and “missing true candidate” cases may change both efficiency and failure modes. Future work should therefore include stress tests with larger candidate sets and more deployment-like shifts, alongside efforts to improve robustness, for instance through self-supervised pretraining on unlabeled spectral data,^4^ and to explore shift-adaptive calibration strategies^22^ when calibration and deployment data are mismatched.

## Associated Content

### Data Availability Statement

The dataset is publicly available at https://github.com/pluskal-lab/MassSpecGym,^1^ and the code for the methods used in this study is available at https://github.com/moranej/ms-cp.

## Supporting information

The Supporting Information is available free of charge:

- SI.pdf: Additional figures and tables. Pearson correlation heatmaps for conditioning variables across scenarios and splits, pairwise distance distributions for conditioning variables, conditional conformal prediction results (MACG and relative set size) for CCCP and CCP-NN across all scenarios, and mean conditional coverage tables for all non-conformity scores and conditioning variables.

## Author Information

### Authors

- **Morteza Rakhshaninejad** – Department of Data Analysis and Mathematical Modeling, Ghent University, Coupure Links 653, Ghent, 9000, Belgium;
- **Gaetan De Waele** – Department of Computer Science, University of Antwerp, Middelheimlaan 1, Antwerp, 2020, Belgium;
- **Mira Jürgens** – Department of Data Analysis and Mathematical Modeling, Ghent University, Coupure Links 653, Ghent, 9000, Belgium;

## Authors’ contributions

M.R., M.J., and W.W. conceptualized the methodology. M.R. designed the study, implemented the framework, conducted all experiments and analyses, and wrote the original draft. G.D.W. developed the retrieval model and training pipeline, provided domain expertise, contributed to methodological discussions, and reviewed the manuscript. M.J. contributed to methodological discussions and reviewed the manuscript. W.W. supervised the project, contributed to methodological discussions, and reviewed the manuscript. All authors read and approved the final version.

## Funding

M.J. and W.W. received funding from the Flemish Government under the “Onderzoeksprogramma Artificiële Intelligentie (AI) Vlaanderen” programme.

## Notes

The authors declare no competing interests.

## Notes

### Competing Interest Statement

The authors have declared no competing interest.

