## Supporting figures and tables for "Reliable Molecular Retrieval from Mass Spectra using Conformal Prediction"

### Supplementary Figures

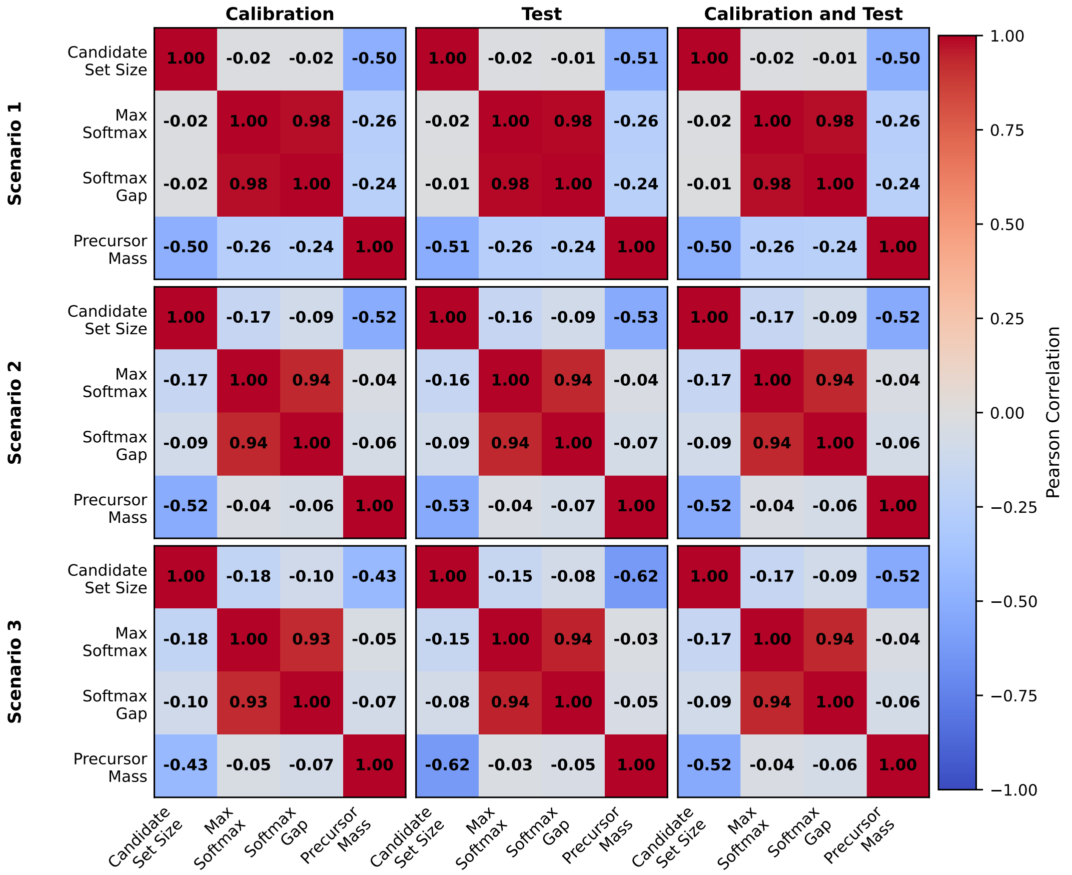


**Figure S1:** Pearson correlation heatmaps for the scalar conditioning variables across scenarios and splits. Rows correspond to Scenario 1–3 and columns correspond to Calibration, Test, and Calibration+Test. Each panel reports Pearson correlation coefficients between Candidate Set Size, Max Softmax, Softmax Gap, and Precursor Mass. Colors encode the correlation strength on a shared scale from −1 to 1 (colorbar at right).

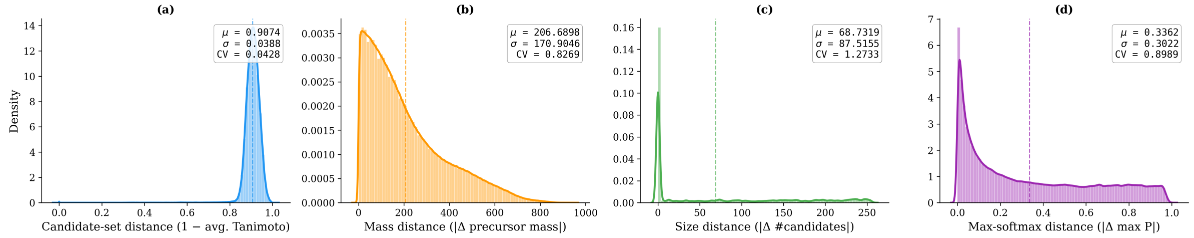


**Figure S2:** Unnormalized pairwise distance distributions for the four conditioning variables. Panels show (a) candidate-set distance defined as $1 - \text{avg. Tanimoto}$, (b) precursor mass distance $|\Delta \text{ precursor mass}|$, (c) candidate set size distance $|\Delta \text{ \#candidates}|$, and (d) max-softmax distance $|\Delta \max P|$. Each panel displays a density histogram with a Gaussian KDE overlay, a dashed vertical line marking the mean, and an inset reporting the mean ($\mu$), standard deviation ($\sigma$), and coefficient of variation (CV).

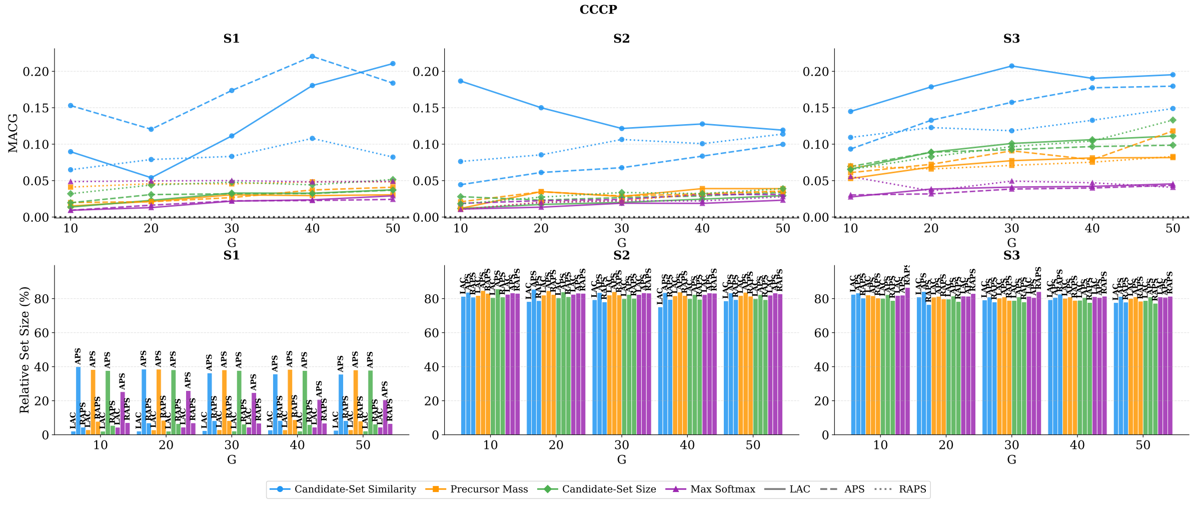


**Figure S3:** Conditional conformal results for CCCP (target coverage 0.9) across Scenarios S1–S3. Top row shows MACG as a function of the number of clusters $G \in \{10, 20, 30, 40, 50\}$. Bottom row shows the mean relative prediction set size $\widehat{R}$ (%) computed from per-sample outputs. Colors and markers indicate the conditioning variable (Candidate-Set Similarity, Precursor Mass, Candidate-Set Size, Max Softmax), and line styles indicate the score function (LAC, APS, RAPS).

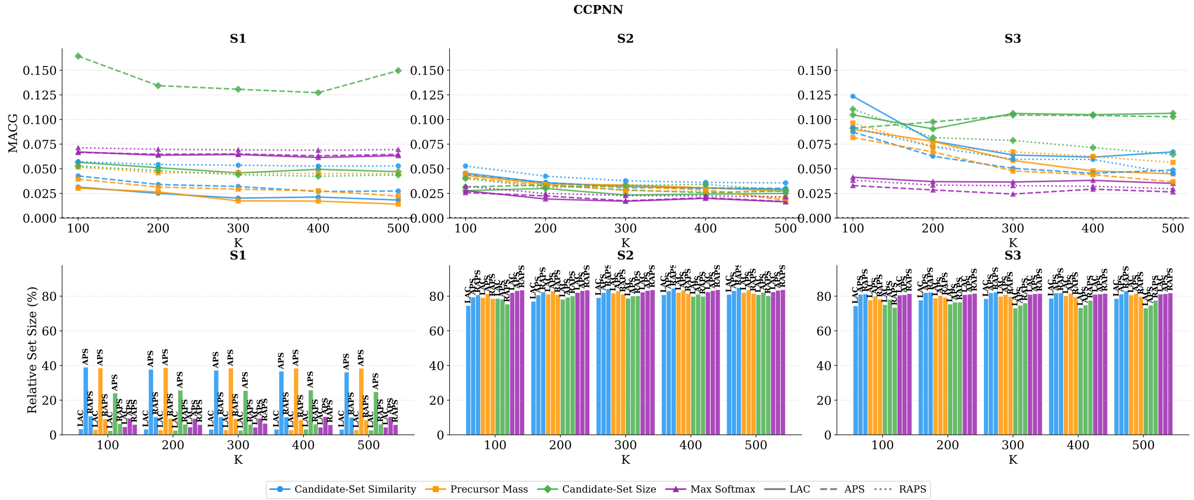


**Figure S4:** Conditional conformal results for CCP-NN (target coverage 0.9) across Scenarios S1–S3. Top row shows MACG as a function of the neighborhood size $K \in \{100, 200, 300, 400, 500\}$. Bottom row shows the mean relative prediction set size $\widehat{R}$ (%) computed from per-sample outputs. Colors and markers indicate the conditioning variable (Candidate-Set Similarity, Precursor Mass, Candidate-Set Size, Max Softmax), and line styles indicate the score function (LAC, APS, RAPS).

### Supplementary Table

**Table S1:** Mean conditional coverage (average of $\widehat{\mathrm{Cov}}_k$ over groups, Eq. (9) in the main text) for all scenarios, non-conformity scores, and conditioning variables at target coverage $1 - \alpha = 0.9$. For CCCP, groups are agglomerative clusters with $G \in \{10, 20, 30, 40, 50\}$. For CCP-NN, groups are $K$-nearest-neighbor neighborhoods with $K \in \{100, 200, 300, 400, 500\}$. Conditioning variables are candidate-set distance (Cand. dist.), precursor mass (Mass), candidate set size (#Cands), and maximum softmax probability (MaxSoftmax).

| | | | CCCP (clusters $G$) | | | | | CCP-NN (neighborhood $K$) | | | | |
|---|---|---|---|---|---|---|---|---|---|---|---|---|
| Scenario | Score | Variable | 10 | 20 | 30 | 40 | 50 | 100 | 200 | 300 | 400 | 500 |
| S1 | LAC | Cand. dist. | 0.911 | 0.912 | 0.854 | 0.790 | 0.765 | 0.906 | 0.910 | 0.910 | 0.912 | 0.911 |
| | | Mass | 0.907 | 0.901 | 0.910 | 0.910 | 0.908 | 0.903 | 0.902 | 0.902 | 0.902 | 0.902 |
| | | #Cands | 0.906 | 0.904 | 0.905 | 0.902 | 0.902 | 0.928 | 0.936 | 0.936 | 0.945 | 0.940 |
| | | MaxSoftmax | 0.901 | 0.902 | 0.901 | 0.903 | 0.901 | 0.959 | 0.959 | 0.958 | 0.959 | 0.960 |
| | APS | Cand. dist. | 0.809 | 0.868 | 0.792 | 0.759 | 0.809 | 0.912 | 0.904 | 0.903 | 0.901 | 0.898 |
| | | Mass | 0.906 | 0.912 | 0.911 | 0.898 | 0.896 | 0.906 | 0.907 | 0.905 | 0.905 | 0.902 |
| | | #Cands | 0.895 | 0.905 | 0.906 | 0.909 | 0.911 | 0.760 | 0.788 | 0.787 | 0.791 | 0.769 |
| | | MaxSoftmax | 0.903 | 0.903 | 0.908 | 0.906 | 0.905 | 0.960 | 0.961 | 0.961 | 0.961 | 0.962 |
| | RAPS | Cand. dist. | 0.946 | 0.961 | 0.971 | 0.953 | 0.981 | 0.956 | 0.954 | 0.953 | 0.952 | 0.951 |
| | | Mass | 0.936 | 0.942 | 0.940 | 0.944 | 0.943 | 0.947 | 0.944 | 0.944 | 0.943 | 0.942 |
| | | #Cands | 0.921 | 0.923 | 0.922 | 0.926 | 0.930 | 0.936 | 0.934 | 0.933 | 0.932 | 0.933 |
| | | MaxSoftmax | 0.949 | 0.949 | 0.949 | 0.946 | 0.945 | 0.969 | 0.968 | 0.969 | 0.968 | 0.970 |
| S2 | LAC | Cand. dist. | 0.745 | 0.790 | 0.812 | 0.806 | 0.829 | 0.889 | 0.893 | 0.898 | 0.902 | 0.902 |
| | | Mass | 0.893 | 0.879 | 0.888 | 0.876 | 0.879 | 0.901 | 0.900 | 0.899 | 0.903 | 0.902 |
| | | #Cands | 0.899 | 0.897 | 0.899 | 0.897 | 0.898 | 0.907 | 0.894 | 0.898 | 0.898 | 0.904 |
| | | MaxSoftmax | 0.901 | 0.901 | 0.897 | 0.899 | 0.896 | 0.902 | 0.901 | 0.901 | 0.901 | 0.901 |
| | APS | Cand. dist. | 0.926 | 0.916 | 0.892 | 0.874 | 0.855 | 0.919 | 0.916 | 0.916 | 0.919 | 0.919 |
| | | Mass | 0.919 | 0.919 | 0.920 | 0.923 | 0.923 | 0.915 | 0.914 | 0.911 | 0.913 | 0.913 |
| | | #Cands | 0.915 | 0.911 | 0.902 | 0.907 | 0.909 | 0.901 | 0.902 | 0.905 | 0.904 | 0.909 |
| | | MaxSoftmax | 0.916 | 0.911 | 0.911 | 0.911 | 0.911 | 0.912 | 0.913 | 0.913 | 0.912 | 0.913 |
| | RAPS | Cand. dist. | 0.855 | 0.857 | 0.845 | 0.851 | 0.836 | 0.930 | 0.931 | 0.932 | 0.930 | 0.931 |
| | | Mass | 0.903 | 0.908 | 0.910 | 0.897 | 0.900 | 0.907 | 0.907 | 0.906 | 0.906 | 0.904 |
| | | #Cands | 0.913 | 0.914 | 0.915 | 0.911 | 0.914 | 0.898 | 0.918 | 0.911 | 0.906 | 0.908 |
| | | MaxSoftmax | 0.901 | 0.899 | 0.900 | 0.901 | 0.897 | 0.917 | 0.916 | 0.916 | 0.915 | 0.916 |
| S3 | LAC | Cand. dist. | 0.809 | 0.781 | 0.741 | 0.765 | 0.771 | 0.820 | 0.847 | 0.854 | 0.857 | 0.853 |
| | | Mass | 0.895 | 0.878 | 0.869 | 0.876 | 0.876 | 0.853 | 0.863 | 0.872 | 0.871 | 0.873 |
| | | #Cands | 0.873 | 0.857 | 0.860 | 0.858 | 0.855 | 0.835 | 0.843 | 0.819 | 0.822 | 0.822 |
| | | MaxSoftmax | 0.893 | 0.884 | 0.886 | 0.883 | 0.882 | 0.872 | 0.874 | 0.874 | 0.874 | 0.876 |
| | APS | Cand. dist. | 0.837 | 0.811 | 0.779 | 0.774 | 0.777 | 0.870 | 0.876 | 0.878 | 0.879 | 0.877 |
| | | Mass | 0.864 | 0.873 | 0.846 | 0.870 | 0.838 | 0.866 | 0.870 | 0.876 | 0.879 | 0.880 |
| | | #Cands | 0.881 | 0.860 | 0.864 | 0.866 | 0.872 | 0.850 | 0.839 | 0.824 | 0.825 | 0.827 |
| | | MaxSoftmax | 0.896 | 0.892 | 0.883 | 0.879 | 0.880 | 0.886 | 0.888 | 0.889 | 0.886 | 0.886 |
| | RAPS | Cand. dist. | 0.865 | 0.865 | 0.855 | 0.839 | 0.819 | 0.870 | 0.883 | 0.889 | 0.884 | 0.900 |
| | | Mass | 0.901 | 0.886 | 0.883 | 0.889 | 0.888 | 0.860 | 0.869 | 0.871 | 0.868 | 0.872 |
| | | #Cands | 0.883 | 0.850 | 0.847 | 0.838 | 0.823 | 0.834 | 0.859 | 0.857 | 0.866 | 0.866 |
| | | MaxSoftmax | 0.949 | 0.917 | 0.923 | 0.907 | 0.900 | 0.889 | 0.890 | 0.889 | 0.890 | 0.891 |
